# Human A2-CAR T cells reject HLA-A2+ human islets transplanted into mice without inducing graft versus host disease

**DOI:** 10.1101/2023.02.23.529741

**Authors:** Cara E. Ellis, Majid Mojibian, Shogo Ida, Vivian C.W. Fung, Søs Skovsø, Emma McIver, Shannon O’Dwyer, Travis D. Webber, Mitchell J.S. Braam, Nelly Saber, Timothy J. Kieffer, Megan K. Levings

**Author notes:** Correspondence information or, Megan Levings: BC Children’s Hospital Research Institute, 950 West 28^th^ Ave, Vancouver BC, V5Z 4H4, Canada. Tim Kieffer: Life Sciences Institute, University of British Columbia, 2350 Health Sciences Mall, Vancouver BC V6T 1Z3, Canada.

## Abstract

**Background:** Type 1 diabetes (T1D) is an autoimmune disease characterised by T cell mediated destruction of pancreatic beta-cells. Islet transplantation is an effective therapy, but its success is limited by islet quality and availability along with the need for immunosuppression. New approaches include use of stem cell-derived insulin-producing cells and immunomodulatory therapies, but a limitation is the paucity of reproducible animal models in which interactions between human immune cells and insulin-producing cells can be studied without the complication of xenogeneic graft-*versus*-host disease (xGVHD).

**Methods:** We expressed an HLA-A2-specific chimeric antigen receptor (A2-CAR) in human CD4+ and CD8+ T cells and tested their ability to reject HLA-A2+ islets transplanted under the kidney capsule or anterior chamber of the eye of immunodeficient mice. T cell engraftment, islet function and xGVHD were assessed longitudinally.

**Results:** The speed and consistency of A2-CAR T cells-mediated islet rejection varied depending on the number of A2-CAR T cells and the absence/presence of co-injected peripheral blood mononuclear cells (PBMCs). When <3 million A2-CAR T cells were injected, co-injection of PBMCs accelerated islet rejection but also induced xGVHD. In the absence of PBMCs, injection of 3 million A2-CAR T cells caused synchronous rejection of A2+ human islets within 1 week and without xGVHD for 12 weeks.

**Conclusions:** Injection of A2-CAR T cells can be used to study rejection of human insulin-producing cells without the complication of xGVHD. The rapidity and synchrony of rejection will facilitate in vivo screening of new therapies designed to improve the success of islet-replacement therapies.

## Introduction

Type 1 diabetes (T1D) is an autoimmune disease characterised by T cell mediated destruction of pancreatic beta-cells. This leads to a loss of insulin production and eventually hyperglycemia^1^. People living with T1D depend on exogenous insulin, via injections or pumps, in combination with glucose monitoring^2^. Transplantation of allogenic islets coupled with immune suppression provides proof of concept that a cell-based therapy can lead to independence from daily insulin injections^3^. However, despite advancements in islet transplant protocols, numerous challenges remain, including limited supply from cadaveric donors, inconsistent islet quality, and the need for long term immunosuppression to mitigate the ongoing risk of allo- and autoimmune rejection^4–6^.

Many groups are working towards increasing the applicability of islet transplantation as a treatment for T1D, focusing on increasing access and reducing the need for immunosuppression. This includes the use of innovative immunomodulatory drugs, tolerogenic immune cell therapies, and encapsulation to prevent physical contact between immune cells and transplanted cells, while allowing transport of nutrients and oxygen to and hormones from the cells^7–9^. There are also substantial efforts to develop islet-replacement strategies through use of pluripotent stem cell-derived insulin-producing cells^10,11^, which can be engineered to evade immune rejection and/or be combined with immunomodulatory therapies. The first clinical trials of genome-edited, pluripotent stem cell-derived insulin producing cells are underway (ClinicalTrials.gov Identifiers: NCT05210530; NCT04786262).

A challenge during the development of these new therapies has been the shortage of animal models for studying and manipulating human immune-cell-driven rejection. These studies typically involve transplantation of islets into immunodeficient mice and engraftment of human peripheral blood mononuclear cells (PBMCs)^12,13 14^. In addition to the varying rate of islet graft rejection in these experimental mice, the onset of xenogeneic graft-*versus*-host disease (xGVHD) limits the window in which graft rejection can be studied without confounding effects^14^. Better animal models in which human cell-mediated islet rejection reliably occurs before onset of xGVHD are needed to increase the feasibility of pre-clinical proof-of-concept studies. One approach is to further genetically modify strains of immunodeficient mice to remove mouse-derived class I and II molecules, which are known T cell targets^15^. Accordingly, NOD-scid IL-2 receptor subunit γ (NSG) mice that lack mouse MHC I and II can be reconstituted with 10-50 million human PBMCs without significant development of xGVHD for >100 days despite ~20% human T cell chimerism in blood. However, these mice are complex to breed, allogeneic islet cell rejection is not synchronous^15^, and it takes up to 10 weeks for rejection of allogeneic insulin-producing stem cells^16^.

We recently created a Chimeric Antigen Receptor (CAR) specific for a common human major histocompatibility antigen, HLA-A2, and showed that it can be used to re-direct the specificity of human Tregs^17^. When expressed in conventional T cells, we speculated that this CAR could be a useful tool to develop a better mouse model of human T cell-mediated islet rejection. Here we report the development and optimization of a method to use human A2-CAR-expressing CD4^+^ and CD8^+^ T cells to mediate rapid and synchronous rejection of human HLA-A2^1^ islets transplanted under the kidney capsule or in the anterior chamber of the eye (ACE) of NSG mice.

## Materials and methods

### A2-CAR T cell generation and analysis

HLA-A2-negative leukopaks were obtained from STEMCELL Technologies (#200-0470) and used according to protocols approved by the University of British Columbia (UBC) Clinical Research Ethics Board. The method to generate A2-CAR-expressing CD4^+^ and CD8^+^ T cells is described^18^. Apoptosis assays were conducted using the Apoptosis/Necrosis Assay kit (Abcam, #ab176749)^18^; cells >60% live, apoptosis-negative were used for injection (**Supplementary Table S1**).

### Animal Studies

All experiments were carried out in accordance with the UBC Animal Care Committee and Canadian Council on Animal Care. Male non-diabetic 8-14 week old NSG mice (005557 NOD.Cg-Prkdc Il2rg/SzJ) were maintained on a 12-h light/dark cycle with ad libitum access to irradiated standard chow diet (#2919) and regular xGVHD monitoring^19^. Blood glucose was measured in saphenous vein samples using a OneTouch Verio glucometer (LifeScan) or in intraperitoneal glucose-stimulated insulin secretion tests (ipGSIS), in which mice were fasted for 6 hours then challenged with 2 g/kg glucose. Human C-peptide and engrafted cell phenotype were measured weekly.

### Human Islet Transplantation

Human cadaveric pancreatic islets **(Supplementary Table S2**) were obtained from the Alberta Diabetes Institute IsletCore and used in accordance with protocols approved by the UBC and Alberta Clinical Research Ethics Boards. For left kidney capsule transplants, each mouse received 2000 IEQ HLA-A2^+^ islets from donor ID R310 or R330. ACE transplants were performed as described^20^; each mouse received 50 HLA-A2 positive or HLA-A2 negative islets from donors R427 or R439. See **SDC Materials and Methods** for more details.

### Human PBMC/T Cell Injection

Mice with islet transplants were randomized based on body weight and stimulated human C-peptide one day or one week before CAR T cell injection. Cryopreserved A2-CAR CD4^+^ and CD8^+^ T cells were thawed with DNAse I solution (STEMCELL) and mixed at equal cell numbers to result in a 50:50 cell ratio suspension in PBS. Autologous PBMCs were thawed in parallel. Cells were injected via the tail-vein.

### Metabolite Analyses

Plasma from the metabolic tests was collected and stored at −30°C until analysis. Human C-peptide was measured from plasma samples using the ALPCO STELLUX Human C-peptide ELISA (80-CPTHU-CH01, Alpco Diagnostics, Salem, NH).

### Statistical Analyses and Data availability

Statistical analyses were performed in R version 4.1.2 ^21^. R code and data available at https://github.com/caraee/A2_CAR. **SDC Materials and Methods.**

## RESULTS

### In the presence of PBMCs, A2-CAR T cells reject HLA-A2^+^ human islets within two weeks

CAR T cells are an effective anti-cancer therapy via their ability to kill tumor cells^22^. To test if they could also be used to reject allogeneic islets, we developed a method to produce T cells expressing an HLA-A2-specific CAR (A2-CAR) (**Supplemental Figure S1**, ^18^). A2-CAR CD4^+^ and CD8^+^ T cells were produced separately to enable injection of defined CD4:CD8 T cell ratios^23,24^.

Human HLA-A2^+^ islets were implanted under the kidney capsule of 14-week-old male, non-diabetic, NSG mice; six mice received sham surgeries. Diabetes was not induced to minimize the risk of reaching a diabetes-related humane endpoint that would prevent long-term monitoring for xGVHD. To assess baseline graft function, three days post-surgery, glucose was injected intraperitoneally to stimulate human C-peptide. Mice were then injected via the tail vein with the indicated number of PBMCs with or without a 1:1 ratio of CD4^+^ to CD8^+^ A2-CAR T cells, such that the total number of injected cells was 10 million. A control group received PBS in the tail vein. Three groups were injected five days post-islet transplantation (PBMC; A2-CAR^lo^-D5; A2-CAR^hi^-D5). To assess how the time of cell injection post-transplant affected rejection, a fourth group was injected 19 days post-islet transplantation (A2-CAR^hi^-D19) (**Figure 1A**).

**Figure 1.**
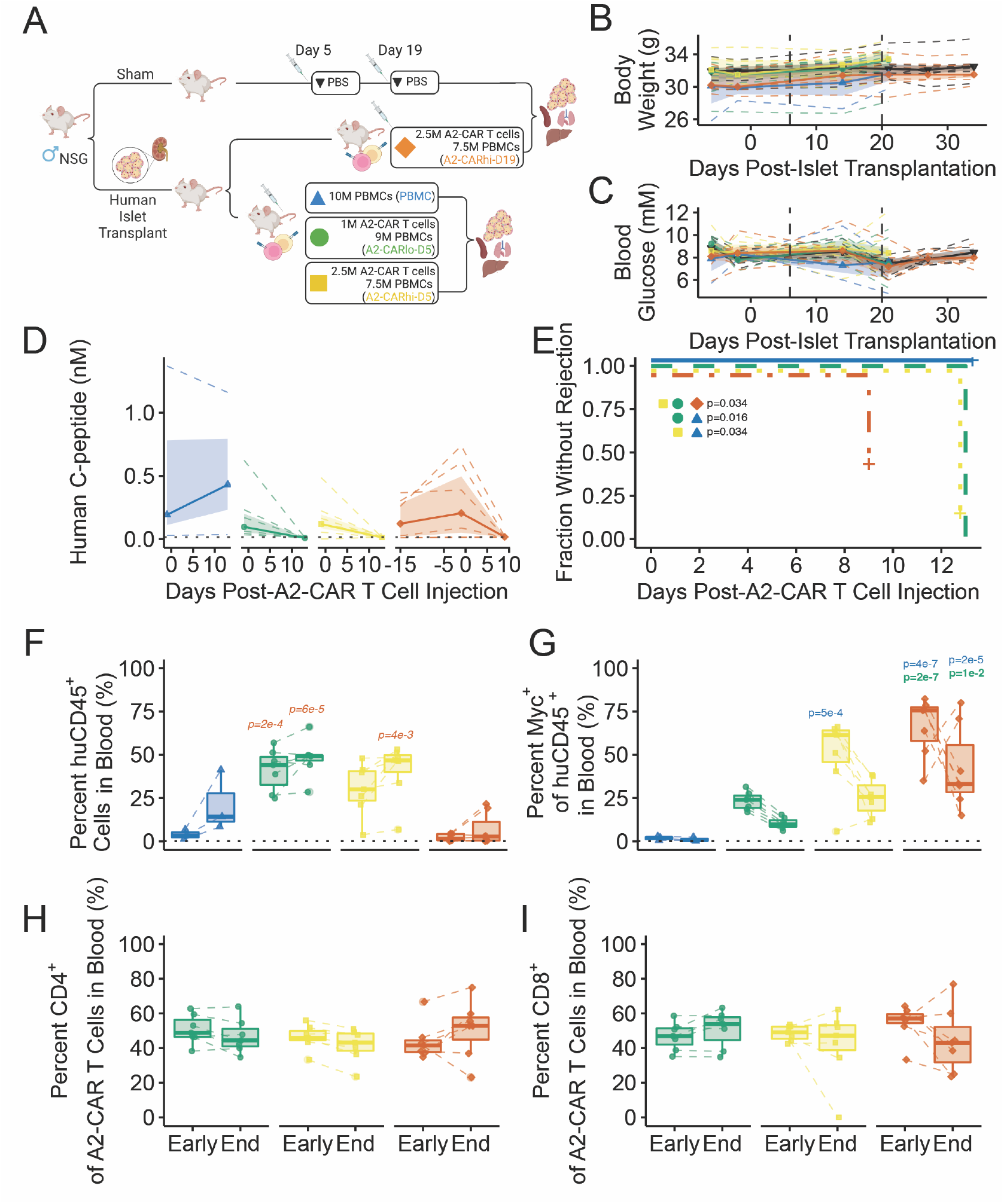
A2-CAR T cells reject A2^+^ islets in the presence of PBMCs. **(A)** Study schematic showing experimental groups and timing of A2-CAR T cell injection. **(B-C)** Body weight and blood glucose were measured after a four hour morning fast. Vertical lines indicate days of CAR-T cell tail vein injections, post-islet transplantation. **(D)** Plasma C-peptide levels collected 30 minutes after glucose delivery (2 g/kg) were measured 1 day before and 9 or 13 days post-A2-CAR T cell injection. **(E)** Survival curve showing the proportion of mice with rejected grafts. **(F-I)** T cell engraftment in blood on day 9 or 13 Early) or 16 (End). The percentages of **(F)** human CD45^+^ and **(G)** A2-CAR (Myc^+^) human CD45^+^ cells were measured in blood. Within the A2-CAR^+^ population, the proportion of **(H)** CD4^+^ and **(I)** CD8^+^ T cells was measured. **(B-D)** Dashed lines are individual mice; solid lines are median values, with the points indicating time of measurements; and shading is the interquartile range. **(E)** Lines indicate fraction of animals without rejection and + indicates censoring. **(F-I)** Horizontal solid lines indicated median, hinges interquartile range (IQR), and whiskers 1.5*IQR. **(D, F, G)** Dotted horizontal lines indicated the limit of detection of human C-peptide **(7.4 pM, D)** or average background of human CD45+ cells in mice in PBS group **(F)**. Colour and style of the p-values indicates direction of comparison: *italics* indicates a comparison to A2-CAR^hi^-D5, regular indicates a comparison to PBMC, and **bold** indicates a comparison to A2-CAR^lo^-D5. **(H,I)** No groups were different from 50%.

Neither islet transplantation nor cell injection impacted body weight (**Figure 1B**) or fasting blood glucose (**Figure 1C**). Six hour fasted plasma human C-peptide levels before CAR T cell injection (**Figure 1D**) ranged from 14.9-1200 pM. There was no evidence for a glucose-stimulated increase in human C-peptide at 30 minutes post-glucose injection pre-CAR-T cells injection, despite a modest increase in blood glucose (**Supplementary Figure S2A**), suggesting that the human islet grafts were not controlling glycemic set point or glucose tolerance^25^. Therefore, we did not use changes in glucose tolerance to assess graft function.

Graft rejection was defined as no detectable human C-peptide at either the fasted or stimulated time points (assay limit of detection: 7.4-14.9 pM after dilution). All mice that received A2-CAR T cells showed an absolute decrease in fasted and stimulated human C-peptide at the first and only time point measured (9 or 13 days) post-CAR-T cell injection (**Figure 1D, Supplementary Figure S2**). Thirteen days after CAR-T cell injection, 7/7 of the A2-CAR^lo^-D5 and 6/7 of the A2-CAR^hi^-D5 mice had rejected grafts whereas none of mice which received PBMCs had rejected grafts (**Figure 1E**). For the A2-CAR^hi^-D19 mice, 4/7 had rejected grafts on day 9 post-A2-CAR T cell injection, but this was not statistically significant compared to PBMC mice (p=0.13); C-peptide was not measured at later time points. Mice that did not meet the criteria for rejection had low but detectable plasma human C-peptide (median: 68 pM, mean: 157 pM, range: 52 to 510 pM), compared to higher human C-peptide levels pre-A2-CAR T cell injection (median: 1800 pM, mean: 1300 pM, range: 40 to 2200 pM) (**Figure 1D, Supplementary Figure S2B**). Human C-peptide was not measured in the blood collected at endpoint (day 16).

Blood was collected to quantify proportions of human CD45^+^ and A2-CAR T cells. All mice received a total of 10 million human immune cells, but those injected with A2-CAR T cells five days post-islet transplantation had higher proportions of human CD45^+^ cells at day 13 (Early) and 16 (End) compared to mice injected with PBMC alone or on day 19 post-islet transplantation (**Figure 1F**), and higher absolute numbers compared to PBMC alone (**Supplementary Figure S3A**; cells were not counted in A2-CAR^hi^-D19 mice). A2-CAR^+^ (Myc^+^) cells had a selective advantage over PBMCs as their proportion and absolute number increased in comparison to the injected A2-CAR^+^/PBMC ratio (**Figure 1G and Supplementary Figure S3B**). A2-CAR^hi^-D19 mice had higher proportions of A2-CAR^+^ compared to A2-CAR^lo^-D5 mice at the Early and End time points, but not compared to the A2-CAR^hi^-D5 mice. Within the A2-CAR^+^ fraction, the ratio of CD4^+^ to CD8^+^ T cells remained stable, with preservation of the 1:1 ratio that was injected at all time points/conditions and no differences between groups (**Figure 1H&I, Supplementary Figure S3C&D**).

Islet grafts were retrieved 16 days post-A2-CAR T cell/PBMC injection. H&E staining of the graft-bearing kidneys showed established islet grafts in all animals except sham surgical mice (**Figure 2A**). Mice that received PBMCs had low-density infiltration of mononuclear cells in all collected tissues (**Figure 2A-C**). In contrast, mice that received A2-CAR T cells had dense infiltration of mononuclear cells around the grafted tissue (**Figure 2D, G, J**), diffuse and sporadic mononuclear cells in kidney and liver (**Figure 2E, H, K**) and denser accumulation in the lung (**Figure 2F, I, L**). No animals showed signs of xGVHD since the study was terminated before the typical onset^15^. However, at the time of islet graft retrieval (16 days post-A2-CAR T cell injection), mice that received 10 million PBMCs had signs of peritoneal inflammation (data not shown).

**Figure 2.**
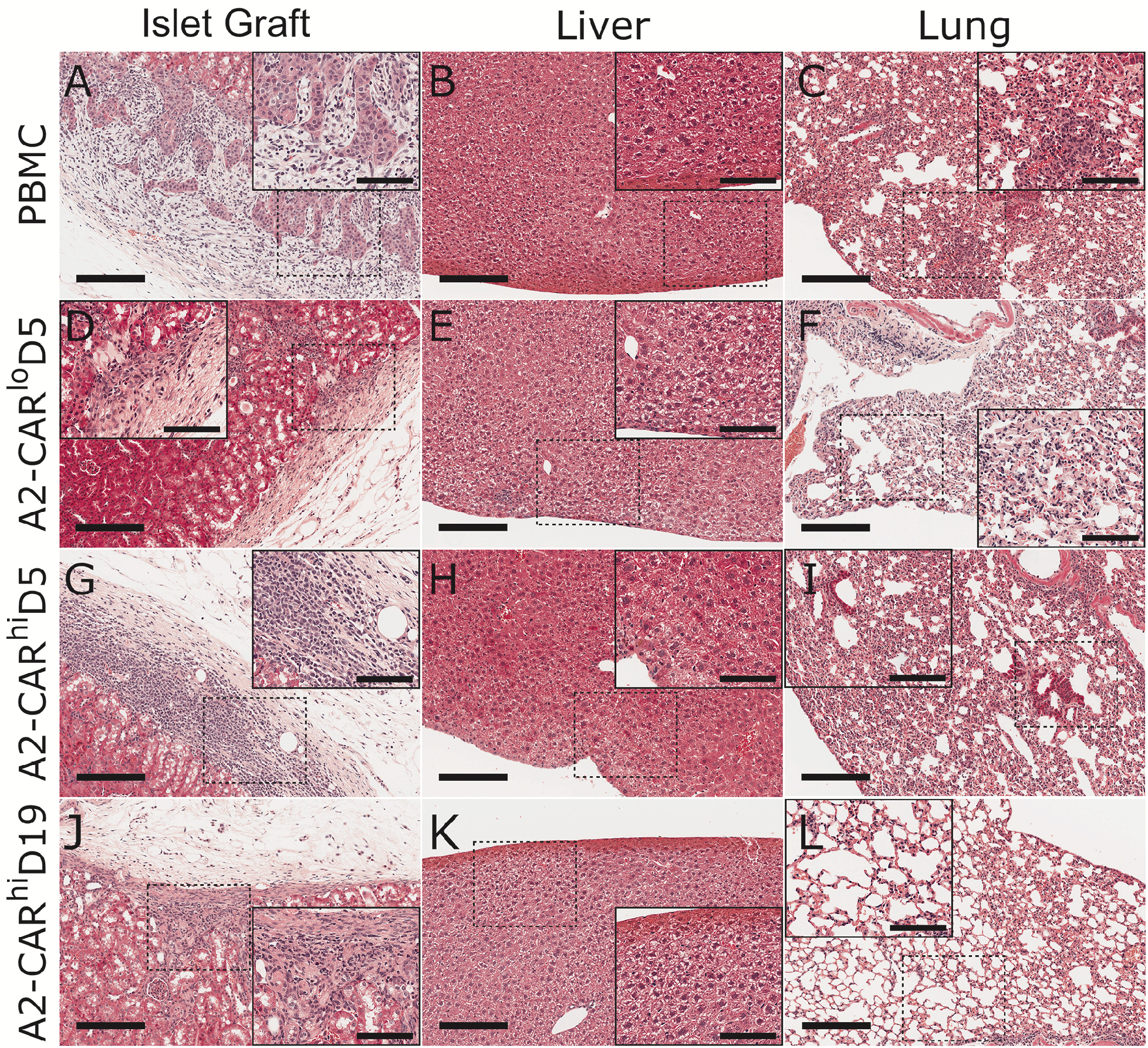
A2-CAR T cell-mediated islet infiltration. Sixteen days post-A2-CAR T cell injection, graft-bearing kidneys **(A, D, G, J)**, livers **(B, E, H, K)** and lungs **(C, F, I, L)** were collected from the indicated groups. Tissue sections were stained with H&E. Scale bars are 200 μm for **A-L** and 100 μm for insets. Images representative of n=7 per group.

### A2-CAR T cell mediated allograft rejection does not require co-injection of PBMCs

We next investigated if PBMC co-injection was necessary to support A2-CAR T cell engraftment and/or allograft rejection; the minimal number of A2-CAR T cells that could induce rejection without xGVHD; and the time course of rejection. Mice transplanted with human HLA-A2^+^ islets under the kidney capsule were injected with 0.5 million A2-CAR T cells (A2-CAR^lo^) or 1.0 million A2-CAR T cells (A2-CAR^med^); A2-CAR T cells in combination with PBMCs (1 million A2-CAR T cells + 2 million PBMCs (A2-CAR^med^PBMC^lo^) or 1 million A2-CAR T cells+ 9M PBMCs, (A2-CAR^med^PBMC^hi^)); or PBS. For mice injected with PBS, a subset (n=4) received 3 million A2-CAR T cells 18 weeks post-islet transplantation (A2-CAR^hi^) (**Figure 3A**).

**Figure 3.**
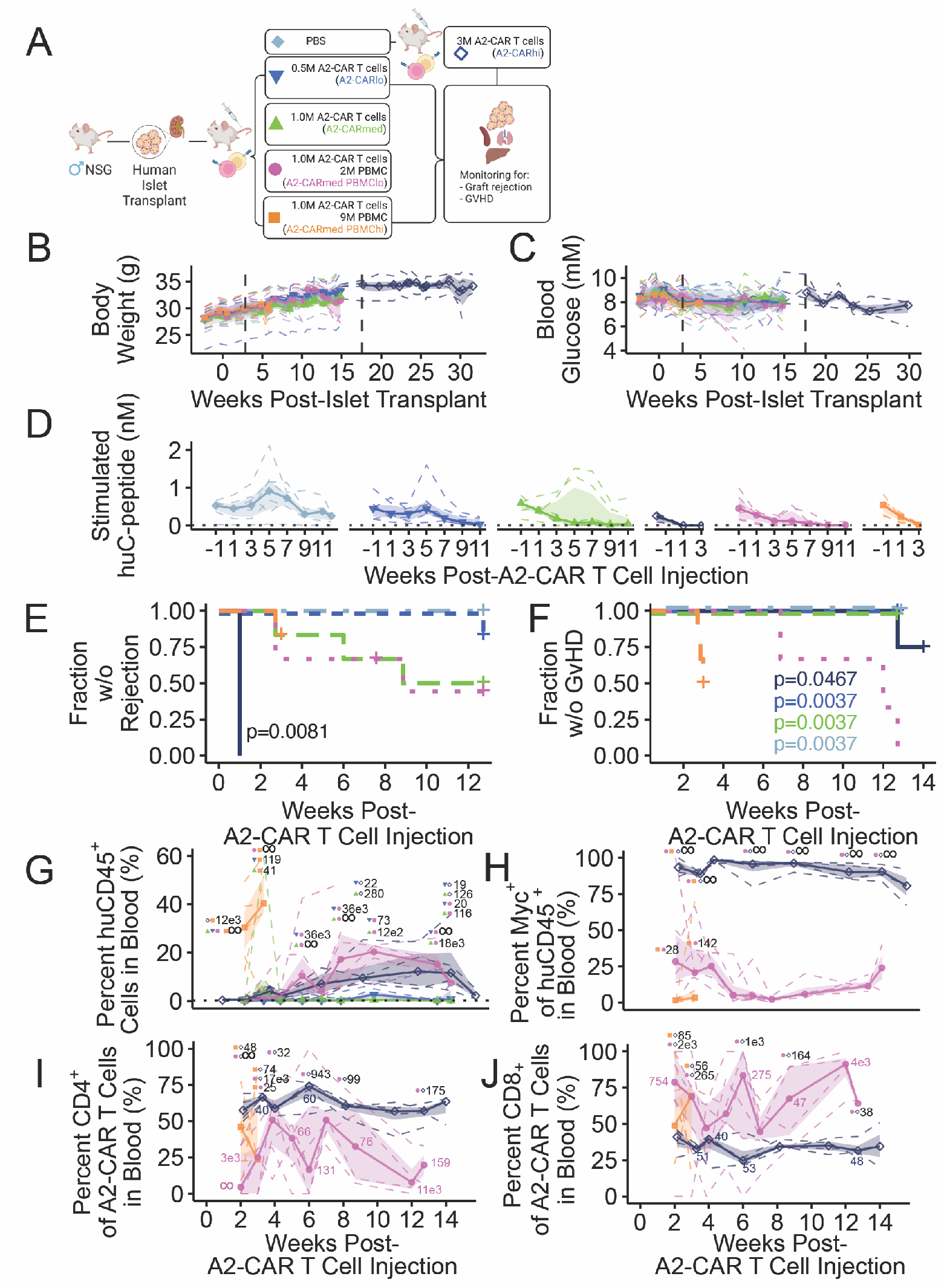
A2-CAR T cells mediate rejection without PBMCs and do not cause xGVHD. **(A)** Study schematic showing experimental groups, timing of islet transplantation, A2-CAR T cell injection, and glucose tolerance tests with blood collection. **(B-C)** Body weights and random blood glucose were measured in the afternoon. Vertical lines indicate days of A2-CAR-T cell injections. **(D)** Mice were challenged with 2 g/kg glucose delivery intraperitoneally after a 6 hour fast, with blood collected at 15 minutes post-glucose injection and assayed for human C-peptide. **(E)** Survival curve showing proportions with rejected grafts. **(F)** Animals were scored for signs of xGVHD. **(G-J)** The proportions of **(G)** human CD45^+^ and **(H)** A2-CAR^+^ (Myc^+^) cells was determined. Within the A2-CAR^+^ cells, the proportions of **(I)** CD4^+^ and **(J)** CD8^+^ cells were measured. **(E, F)** Lines indicate the fraction of animals without rejection/xGVHD, with + indicating animals that were euthanized without reaching the criteria. The A2-CAR^hi^ mice were four of the six mice that initially received only PBS. In all other panels, solid lines are the medians with points indicating the times of measurements, shading is the interquartile range, and dashed lines are individual mice. **(D, G)** Dotted horizontal lines indicated the limit of detection of human C-peptide **(14.9 pM, D)** or average background of human CD45+ cells in mice in the PBS group **(G)**. **(G-J)** All values are evidence ratios, indicating the number of times more likely it is that a difference is observed compared to no difference. Values next to symbols show the evidence ratio for comparisons between those groups. **(I-J)** Values without symbols show the evidence ratio for comparison to the injected ratio, 50%. If no result is reported, the posterior probability did not exceed 95% and/or the absolute difference was less than 10% (low biological significance).

Transplantation and A2-CAR T cell injection did not impact body weight (**Figure 3B**), or random fed (**Figure 3C**) or six hour fasted blood glucose (**Supplementary Figure S4A**). Human C-peptide and intraperitoneal glucose tolerance were measured one week before A2-CAR T cell injection, then every other week to a maximum of 11 weeks post-injection (**Figure 3D**). Six hour fasted plasma human C-peptide levels were variable before A2-CAR T cell injection, ranging from 20 pM to 1200 pM (**Supplementary Figure S4B**), and as in **Figure 1** there was no evidence for a glucose-stimulated increase in human C-peptide (**Supplementary Figure S4A**). PBS mice had stable stimulated human C-peptide throughout the study, and no mice met the criteria for graft rejection. For mice injected with A2-CAR T cells alone, there was a dosedependent effect on C-peptide decline: A2-CAR^hi^ mice had a rapid and consistent human C-peptide decline whereas A2-CAR^lo^ and A2-CAR^med^ mice had a more gradual decline. For mice receiving a mixture of A2-CAR T cells and PBMCs: 1/6 A2-CAR^med^PBMC^hi^ had rejected grafts at three weeks post-A2-CAR T cell injection and the group was terminated due to the rapid development of xGVHD (**Figure 3E&F**). Three of six A2-CAR^med^PBMC^lo^ mice had rejected grafts 13 weeks post-CAR T cell injection.

For mice only receiving A2-CAR T cells without PBMCs, at week 13 post-A2-CAR-T cell injection 1/6 A2-CAR^lo^ and 3/6 A2-CAR^med^ mice met the criteria for rejection (**Figure 3E**). In contrast, at one week post-A2-CAR T cell injection, 4/4 A2-CAR^hi^ mice had undetectable plasma human C-peptide, resulting in greater rejection compared to all other groups. Several mice had decreased but detectable human C-peptide (**Figure 3D, Supplementary Figure S4B**) so they may have progressed to rejection had the study been longer (+ symbol indicates animals that were euthanized before meeting the criteria for rejection).

xGVHD was assessed multiple times per week and prior to week 12 was exclusively observed in mice that received PBMCs. Three of six A2-CAR^med^PBMC^hi^ mice developed severe xGVHD requiring euthanasia three weeks post cell injection, whereas A2-CAR^med^PBMC^lo^ mice began developing xGVHD seven weeks post cell injection. Only one of four A2-CAR^hi^ mice, but no A2-CAR^lo^ or A2-CAR^med^ mice, showed signs of xGvHD at the time of tissue collection (14 weeks post cell injection). A2-CAR^med^PBMC^lo^ mice had greater development of xGVHD compared to PBS, A2-CAR^lo^, A2-CAR^med^ and A2-CAR^hi^ mice (**Figure 3F**). No other groups were different, but many animals were removed from the study before they met the criteria for xGVHD (+ symbol in Figure 3F), diminishing our ability to detect statistically significant differences.

Blood was collected weekly to measure human immune cell engraftment and random fed human C-peptide (**Figure 3G-J**, **Supplementary Figure S5**). Injection of PBMCs resulted in higher percentages (**Figure 3G, Supplementary Figure S6A**) and numbers (**Supplementary Figure S7A**) of human CD45+ cells in blood compared to CAR T cells alone. A2-CAR^med^PBMC^hi^ mice had higher percentages of human CD45^+^ cells in their blood compared to A2-CAR^med^PBMC^lo^ mice or A2-CAR^hi^ mice at two weeks post-A2-CAR T cell injection, as well as compared to A2-CAR^lo^ mice or A2-CAR^med^ at two and three weeks (**Figure 3G**, **Supplementary Figure S7A**). Similarly, A2-CAR^med^PBMC^lo^ mice had more human CD45^+^ cells compared to A2-CAR^lo^ mice or A2-CAR^med^ mice at all points measured from four until 13 weeks post-A2-CAR T cell injection. In concordance with the increased dose, A2-CAR^hi^ mice had higher percentages of human CD45^+^ cells in their blood compared to A2-CAR^lo^ mice or A2-CAR^med^ mice at all points measured from four until 13 weeks post-A2-CAR T cell injection (cell numbers were not measured in the A2-CAR^hi^ mice).

Mice injected with a higher dose of A2-CAR T cells had higher proportions of Myc^+^ A2-CAR cells within human CD45^+^ cells (**Figure 3H**, **Supplementary Figure S7B**). A2-CAR^lo^ and A2-CAR^med^ mice had very few (<1%) human CD45^+^ cells in their blood so these groups were not analyzed further. A2-CAR^hi^ mice had higher proportions of Myc^+^ cells at all time points measured compared to A2-CAR^med^PBMC^lo^ mice or A2-CAR^med^PBMC^hi^ mice. A2-CAR^med^PBMC^lo^ mice had moderately higher percentages of human CD45^+^ cells that were also positive for Myc compared to A2-CAR^med^PBMC^hi^ mice, reflecting the higher number of CD45^+^ PBMCs in the A2-CAR^med^PBMC^hi^ mice and thus the lower relative engraftment.

We also measured the percentages of CD4^+^ (**Figure 3I, Supplemental Figure S6C**) and CD8^+^ (**Figure 3J**, **Supplemental Figure S6D**) T cells within A2-CAR^+^ cells, and numbers in all groups but A2-CAR^hi^ (**Supplemental Figure S7C&D**). A2-CAR^hi^ mice had higher percentages of CD4^+^ A2-CAR T cells, and corresponding lower percentages of CD8^+^ A2-CAR T cells, compared to A2-CAR^med^PBMC^lo^ mice or A2-CAR^med^PBMC^hi^ mice. A2-CAR^med^PBMC^hi^ mice had higher percentages of CD4^+^ A2-CAR T cells at two weeks post-cell injection compared to A2-CAR^med^PBMC^hi^ mice, but we did not observe evidence for any differences in CD8^+^ A2-CAR T cells between these groups. The percentages of CD4+ and CD8+ A2-CAR T cells were not stable over time in the A2-CAR^med^PBMC^lo^ mice and differed from the 1:1 injection ratio, likely because the absolute numbers of CAR^+^ T cells were low (A2-CAR^med^PBMC^lo^ median: 94 cells/mL; mean: 295 cells/mL; range 0-2560 cells/mL, **Supplementary Figure S6B**), so a small fluctuation in number of cells detected translates to a larger fluctuation in percentage. The ratio of CD4^+^:CD8^+^ A2-CAR T cells was more stable, with a possible selective advantage for CD4^+^ A2-CAR T cells.

### A2-CAR T cells home to grafts, while PBMCs infiltrate other tissues

Grafts were retrieved post-A2-CAR T cell injection: after three weeks for the A2-CAR^med^PBMC^hi^ recipients and 12-14 weeks for the remaining groups. A2-CAR^lo^ mice had visible islet grafts in 5/6 animals, correlating with 5/6 having detectable human C-peptide at the time of graft collection. Two grafts showed signs of immune cell infiltration, with no other tissues showing signs of infiltration (**Figure 4**). Five of six A2-CAR^med^ mice had identifiable human islet grafts with signs of infiltration and fibrosis, while the graft from the remaining mouse was not visible. Three of the six mice in this group had detectable C-peptide (**Figure 3D-E)**. Similarly to A2-CAR^lo^ mice, no other tissues showed signs of infiltration for any of the A2-CAR^med^ mice. A2-CAR^hi^ mice had identifiable human islet grafts with infiltration, fibrosis, and no visible endocrine tissue; although only one mouse had signs of xGVHD at the time of graft collection, all livers and lungs had signs of immune cell infiltration. Control PBS mice had clearly visible human islet grafts with no signs of infiltration in the livers or lungs. All visible grafts from A2-CAR^med^PBMC^lo^ and A2-CAR^med^PBMC^hi^ mice, had clear signs of infiltration and infiltrating cells in 6/6 livers and lungs.

**Figure 4:**
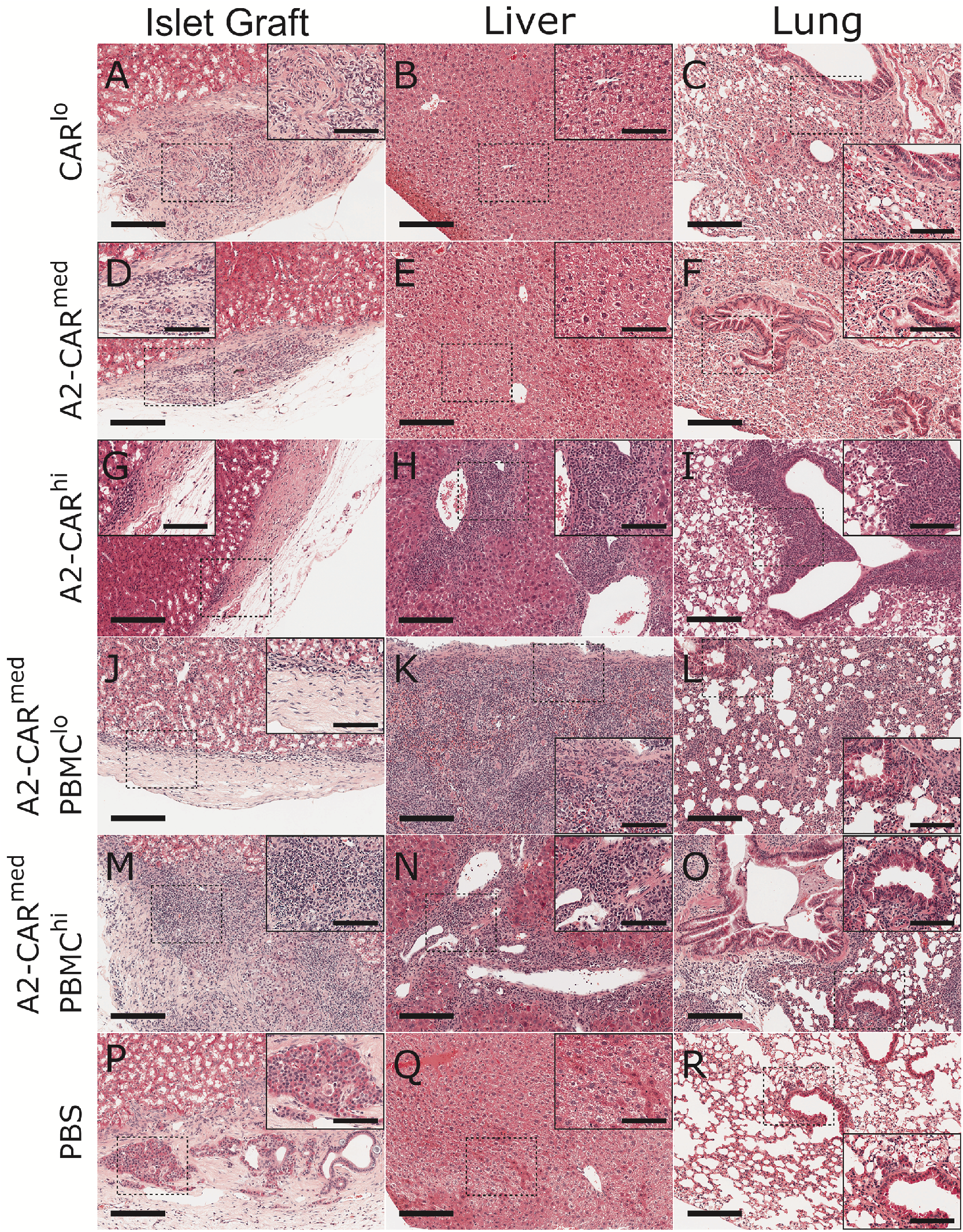
A2-CAR T cells specifically home to the graft and destroy human islets. Graft-bearing kidneys **(A, D, G, J, M, and P)**, livers **(B, E, H, K, N, and Q)** and lungs **(C, F, I, L, O, and R)** were collected and tissue sections were stained with H&E. Scale bars are 200 μm for A-L and 100 μm for insets. Images representative of n=6 **(A-F, J-O)**, 4 **(G-I)**, or 2 **(P-R)**.

### A2-CAR T cell-mediated rejection of A2^+^ human islets in the anterior chamber of the eye

Although islet transplantation in mice is typically done under the kidney capsule, alternate sites such as the spleen and anterior chamber of the eye have advantages such as rapid vascularization and the ability for longitudinal graft imaging^20,26^. To assess if A2-CAR T cells could also mediate efficient islet graft rejection in an alternate site, human HLA-A2^pos^ or HLA-A2^neg^ islets from two different donors were transplanted into the anterior chamber of the eye (ACE, **Figure 5A**). Once islet grafts were vascularized (as evaluated by observation under a stereomicroscope), mice were injected with the optimal cell dose for rapid rejection identified in **Figure 3**: 3 million A2-CAR T cells at a 1:1 ratio of CD4^+^ to CD8^+^.

**Figure 5:**
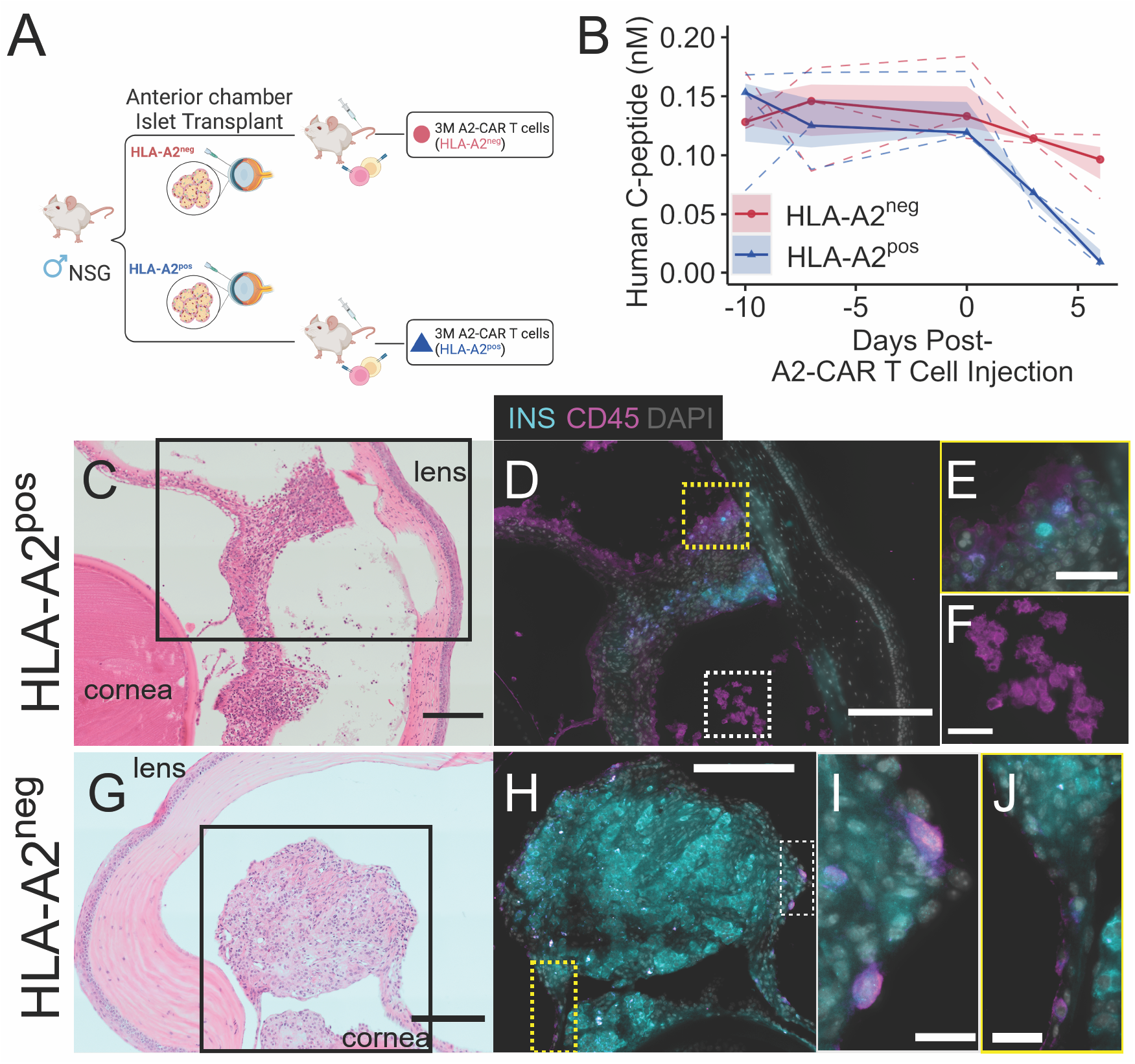
Antigen-specific rejection of islets transplanted in the anterior chamber. **(A)** Study schematic showing experimental groups, timing of islet transplantation, A2-CAR T cell injection, and blood collection. **(B)** Following a 4 hour fast, mice were provided food for 45 minutes then blood was collected and human C-peptide was measured in plasma. Graft-bearing eyes were retrieved from 6 days post-A2-CAR T cell injection and stained for **(C, G)** H&E and **(D-J)** insulin (cyan), and human CD45 (magenta). In B, points and solid lines are the medians, shading is the interquartile range, and dashed lines are individual mice. Scale bars are 200 μm **(C, D, G, H)**, 40 μm **(E, F)**, or 25 μm **(I, J)**. Images representative of n=3.

On day six post T cell injection, mice with HLA-A2^pos^ islet transplants had lower human C-peptide levels compared to all time points pre-A2-CAR T cell injection (**Figure 5B**). In contrast, mice transplanted with HLA-A2^neg^ islets did not have a decrease in human C-peptide (greatest difference was observed on day 6 compared to day 0, p=0.58, 50±20 pM lower). There were minimal differences in human C-peptide observed between the HLA-A2^pos^ and HLA-A2^neg^ mice at any time points measured (at day 6 HLA-A2^pos^ mice had 80±20 pM lower C-peptide (p=0.16)).

In support of the human C-peptide data, islet grafts from mice with HLA-A2^pos^ islets had signs of immune cell infiltration in the islet graft (**Figure 5C**), with minimal insulin immunoreactivity and many human CD45^+^ immune cells within and surrounding the graft (**Figure 5D-E).** In contrast, islet grafts from mice with HLA-A2^neg^ islets had minimal signs of immune cell infiltration and strong insulin immunoreactivity (**Figure 5G-J**).

## DISCUSSION

Here, we show that human T cells engineered with HLA-specific CARs can be leveraged as a pre-clinical mouse model of allograft rejection without the complication of xGVHD. Injection of 3 million A2-CAR T cells into graft-bearing mice caused rapid, synchronous and antigen-specific rejection of human islets transplanted in two different sites. This model can be leveraged to test a variety of therapeutic modalities for T1D.

In pursuit of a model that circumvented xGVHD, we compared the efficacy of A2-CAR T cells in the presence or absence of co-injected autologous PBMCs. We found that co-injection of PBMCs and A2-CAR T cells increased human cell engraftment but was not necessary for engraftment or rejection. In the system reported here, injection of 3 million A2-CAR T cells without PBMCs provided consistent engraftment and rejection within 1 week without the complication of xGVHD for at least 12 weeks. The rapidity of rejection and lack of requirement for PBMCs is similar to data reported by Muller et al. who injected 2 million A2-CAR T cells and observed rejection of A2-transgenic islets within 1-2 weeks^26^. Lower doses of A2-CAR T cells without PBMCs also induced rejection without xGVHD, but at a delayed rate, with half of the mice receiving 1 million A2-CAR T cells rejecting grafts by 13 weeks post-injection.

Models involving implantation of insulin-producing cells from stem cells often require extensive time to mature in vivo^10^. Previous data suggest that the time between transplant and injection of human cells could affect rejection kinetics^14^, possibly due to lower surgery-related inflammation. Thus, we compared the effectiveness of A2-CAR T cell mediated rejection upon delayed injection 19 days post-transplant. Despite this delay, we observed graft rejection within one week, indicating that recently-transplanted and stable grafts are similarly susceptible to A2-CAR T cell-mediated rejection. Human CD45^+^ cell engraftment was lower in mice injected on day 19 compared to those injected on day five post-transplant, but the probability of graft rejection was similar, likely due to efficient homing to the graft^26,27^. When we injected a higher dose of A2-CAR-T cells without PBMCs 18 weeks post-islet transplantation, we observed rapid and consistent rejection despite low human CD45^+^ cell engraftment, indicating that this system would allow testing of a variety of immune evasion and immunomodulatory strategies in stem cell-derived insulin-producing grafts even after a maturation period.

The clinical production of CAR-T cells commonly involves transduction of unselected PBMCs, leading to variation in the proportion of transduced CD4^+^ and CD8^+^ T cells^23^. To decrease donor-to-donor variability, we pre-selected CD4^+^ and CD8^+^ T cells so that defined proportions of purified CAR^+^ cells could be used for injection. We found that the injected ratio remained relatively constant over time, with persistence of both CD4^+^ and CD8^+^ T cells. In contrast, with PBMC-mediated rejection, the CD4^+^:CD8^+^ T cell ratio fluctuates over time, with preferential expansion of CD8^+^ T cells in mice with human islet allografts^15^. The ability to simultaneously study CD4^+^ and CD8^+^ T cell rejection in the A2-CAR T cell-rejection model could thus have advantages in terms of testing the efficacy of various immune evasion/tolerance strategies.

In conclusion, here we investigated the ability of alloantigen-specific A2-CAR-T cells to mediate rejection of human islets transplanted in two different locations in NSG mice. The ability of these cells to swiftly mediate rejection without the risk of xGVHD sets the stage to use this model to test the effectiveness of immune-evasion and/or tolerance induction strategies for T1D.

## AUTHORSHIP PAGE

### Author Contributions

**CE** research design, paper writing, performance of the research, data analysis, statistical analyses

**MM** research design, paper writing performance of the research, data analysis

**SI** research design, performance of the research

**VF** research design, paper writing performance of the research, data analysis

**SS** paper writing, data analysis, study schematics, data analysis, statistical analyses

**EM** performance of the research

**SOD** research design, performance of the research

**TW** research design, performance of the research

**MB** performance of the research

**NS** performance of the research

**TJK** research design, paper writing

**MKL** research design, paper writing

### Disclosures

The authors received funding from CRISPR Therapeutics for a portion of this work. At the time of submission, TJK was employed by ViaCyte, Inc.

### Funding

CRISPR Therapeutics, JDRF Canada (3-COE-2022-1103-M-B) and the National Institutes of Health (5R01DK120392-03). SI was supported by a fellowship from the Manpei Suzuki Diabetes Foundation. MKL receives a salary award from the BC Children’s Hospital Research Institute and holds a Tier 1 Canada Research Chair in Engineered Immune Tolerance.

## ABBREVATIONS

A2-CAR: HLA-A2 expressing Chimeric antigen receptor
ACE: anterior chamber of the eye
BF: Bayes factor
CCV: cytocalcein violet
CMRL: Connaught medical research laboratorie
DNase: deoxyribonuclease
HLA-A2: human leukocyte antigen receptor A gene serotype 2
IL-2: interleukin 2
IPGTT: intraperitoneal glucose tolerance test
MHC: major histocompatibility class
NGFR: nerve growth factor receptor
NOD: non-obese diabetic
NSAID: non-steroidal anti-inflammatory drugs
NSG: NOD scid gamma
PBMCs: peripheral blood mononuclear cells
PBS: phosphate buffered saline
PFA: paraformaldehyde
RBC: red blood cells
T1D: type 1 diabetes
xGVHD: xenogeneic graft-*versus*-host disease

## Acknowledgements

We thank the Alberta Diabetes Institute IsletCore program for their work in procuring human donor pancreases for research. We especially thank the organ donors and their families for their generous and meaningful gifts in support of research. We thank Dr. John Dimitry and Mr. Peter Overby with their assistance in improving accessibility of the figures. The University of British Columbia is situated on unceded Indigenous territories, including the territories of the S<u>k</u>w<u>x</u>wú7mesh (Squamish), səĺilwətaɁɬ (Tsleil-Waututh), and x^w^məθk^w^θýθm (Musqueam) First Nations.

## References

1. Atkinson MA, Eisenbarth GS, Michels AW. Type 1 diabetes. Lancet. 2014;383(9911): 69–82.

2. Domingo-Lopez DA, Lattanzi G, L HJS, et al. Medical devices, smart drug delivery, wearables and technology for the treatment of Diabetes Mellitus. Adv Drug Deliv Rev. 2022;185: 114280.

3. Pepper AR, Bruni A, Shapiro AMJ. Clinical islet transplantation: is the future finally now? Curr Opin Organ Transplant. 2018;23(4): 428–439.

4. Walker S, Appari M, Forbes S. Considerations and challenges of islet transplantation and future therapies on the horizon. Am J Physiol Endocrinol Metab. 2022;322(2): e109–E117.

5. Shapiro AM, Pokrywczynska M, Ricordi C. Clinical pancreatic islet transplantation. Nat Rev Endocrinol. 2017;13(5): 268–277.

6. Hart NJ, Powers AC. Use of human islets to understand islet biology and diabetes: progress, challenges and suggestions. Diabetologia. 2019;62(2): 212–222.

7. Marfil-Garza BA, Polishevska K, Pepper AR, Korbutt GS. Current State and Evidence of Cellular Encapsulation Strategies in Type 1 Diabetes. Compr Physiol. 2020;10(3): 839–878.

8. Pathak S, Meyer EH. Tregs and Mixed Chimerism as Approaches for Tolerance Induction in Islet Transplantation. Front Immunol. 2020;11: 612737.

9. Raffin C, Vo LT, Bluestone JA. Treg cell-based therapies: challenges and perspectives. Nat Rev Immunol. 2020;20(3): 158–172.

10. Ellis C, Ramzy A, Kieffer TJ. Regenerative medicine and cell-based approaches to restore pancreatic function. Nat Rev Gastroenterol Hepatol. 2017;14(10): 612–628.

11. Verhoeff K, Cuesta-Gomez N, Jasra I, Marfil-Garza B, Dadheech N, Shapiro AMJ. Optimizing Generation of Stem Cell-Derived Islet Cells. Stem Cell Rev Rep. 2022.

12. Kenney LL, Shultz LD, Greiner DL, Brehm MA. Humanized Mouse Models for Transplant Immunology. Am J Transplant. 2016;16(2): 389–397.

13. Kooreman NG, de Almeida PE, Stack JP, et al. Alloimmune Responses of Humanized Mice to Human Pluripotent Stem Cell Therapeutics. Cell Rep. 2017;20(8): 1978–1990.

14. King MA, Covassin L, Brehm MA, et al. Human peripheral blood leucocyte non-obese diabetic-severe combined immunodeficiency interleukin-2 receptor gamma chain gene mouse model of xenogeneic graft-versus-host-like disease and the role of host major histocompatibility complex. Clin Exp Immunol. 2009;157(1): 104–118.

15. Brehm MA, Kenney LL, Wiles MV, et al. Lack of acute xenogeneic graft-versus-host disease, but retention of T-cell function following engraftment of human peripheral blood mononuclear cells in NSG mice deficient in MHC class I and II expression. FASEB J.2019;33(3): 3137–3151.

16. Sintov E, Nikolskiy I, Barrera V, et al. Whole-genome CRISPR screening identifies genetic manipulations to reduce immune rejection of stem cell-derived islets. Stem Cell Reports.2022;17(9): 1976–1990.

17. MacDonald KG, Hoeppli RE, Huang Q, et al. Alloantigen-specific regulatory T cells generated with a chimeric antigen receptor. J Clin Invest. 2016;126(4): 1413–1424.

18. Fung VCW, Rosado-Sanchez I, Levings MK. Transduction of Human T Cell Subsets with Lentivirus. Methods Mol Biol. 2021;2285: 227–254.

19. Dawson NAJ, Rosado-Sanchez I, Novakovsky GE, et al. Functional effects of chimeric antigen receptor co-receptor signaling domains in human regulatory T cells. Sci Transl Med. 2020;12(557).

20. Mojibian M, Harder B, Hurlburt A, Bruin JE, Asadi A, Kieffer TJ. Implanted islets in the anterior chamber of the eye are prone to autoimmune attack in a mouse model of diabetes. Diabetologia. 2013;56(10): 2213–2221.

21. R: A language and environment for statistical computing [computer program]. Vienna2022.

22. June CH, Sadelain M. Chimeric Antigen Receptor Therapy. N Engl J Med. 2018;379(1): 64-73.

23. Turtle CJ, Hanafi LA, Berger C, et al. CD19 CAR-T cells of defined CD4+:CD8+ composition in adult B cell ALL patients. J Clin Invest. 2016;126(6): 2123–2138.

24. Sommermeyer D, Hudecek M, Kosasih PL, et al. Chimeric antigen receptor-modified T cells derived from defined CD8+ and CD4+ subsets confer superior antitumor reactivity in vivo. Leukemia. 2016;30(2): 492–500.

25. Rodriguez-Diaz R, Molano RD, Weitz JR, et al. Paracrine Interactions within the Pancreatic Islet Determine the Glycemic Set Point. Cell Metab. 2018;27(3): 549–558 e544.

26. Muller YD, Ferreira LMR, Ronin E, et al. Precision Engineering of an Anti-HLA-A2 Chimeric Antigen Receptor in Regulatory T Cells for Transplant Immune Tolerance. Front Immunol.2021;12: 686439.

27. Dawson NA, Lamarche C, Hoeppli RE, et al. Systematic testing and specificity mapping of alloantigen-specific chimeric antigen receptors in regulatory T cells. JCI Insight. 2019;4(6).

