## Supplemental Data for "Human A2-CAR T cells reject HLA-A2+ human islets transplanted into mice without inducing graft versus host disease"

### SDC Materials and Methods

#### Human Islet Culture

Islets were incubated at 37°C/5% $CO_2$ in supplemented CMRL media at a density of 1000 IEQ per 10 mL in 100 mm x 20 mm non-tissue culture treated petri dishes (Fisher Scientific FB0875712) for 2-5 days before transplantation. CMRL media (Corning 98-304-CV) was obtained from the Alberta Diabetes Institute, and made according to the following protocol: 500 mL of CMRL, 5 mL ITS (Corning 25-800-CR), 5 mL glutaMAX (Gibco 35050-061), 8.5 mL 30% BSA (final 0.5%) (Equitech Bio, BL64-1000), 2.5 mL Pen/strep (Lonza 09-757F).

#### Human Islet Transplantation under the kidney capsule

Mice were randomized based on body weight measured the day before islet transplantation, matching median and IQR within 10%. Mice were anaesthetised with inhalable isoflurane and given 5 mg/kg Anafen, 8 mg/kg Bupivacaine, 0.05 mg/kg buprenorphine, 10 mg/kg Baytril, and 200 µL Lactate Ringer's Saline subcutaneously. Each mouse received 2000 IEQ HLA-A2$^+$ human islets from donor ID R310 (**Figure 1-2**) or R330 (**Figure 3-4**) (**Supplementary Table S2**). All procedures were performed under the sterile tip method, where the tips of instruments were heated to 300°C using a glass bead sterilizer between mice instead of new instruments each time. Mice were given HydroGel on the cage bottom post-surgically and monitored for recovery. Grafts were retrieved from mice in a terminal procedure 16 days post-transplant (**Figures 1-2**); or upon reaching human endpoint due to xGvHD, or 12-14 weeks post-CAR-T cell injection (**Figure 3-4**).

#### Human Islet Transplantation into the Anterior Chamber of the Eye

For anterior chamber of the eye (ACE) transplants, each mouse received 50 HLA-A2 positive or HLA-A2 negative islets from donors R427 and R439 (**Supplementary Table S2**) into the anterior chamber of the eye. Mice were injected subcutaneously with a mixture of the analgesic NSAID Metacam (5 mg/kg) and Buprenorphine (0.05 mg/kg) 15-20 minutes before surgery started. The head of the mice was fixed in a head holder equipped with a nose cone, positioned under a stereomicroscope (Nikon, SMZ-745T). One drop of alcaine (0.5%) (DIN 00035076; Novartis Pharmaceuticals, Canada), a local anesthetic, and Refresh Tears (DIN 02231008) supplemented with 0.3% gentamicin (wt/vol), was applied to the eye 5 minutes prior to islet injection. The mice were further anesthetised with inhalable isoflurane via a nose cone (2-5%). The cornea was punctured obliquely using a 27G needle, and the needle was passed into the anterior chamber close to the sclera, avoiding damage to the iris, lens or bleeding. Islets were transplanted into the anterior chamber of the eye using a stripper micropipette (MXL3-BP-IND-200; Origio MidAtlantic Devices, Mt Laurel, NJ, USA) and a microinjector (Mitutoyyo) attached to the modified micromanipulator via polyethylene-50 tubing (#427517, BD Clay Adams. The site of entry was widened with heat-sterilised fine forceps, the micropipette containing the islets was passed into the anterior chamber and islets were ejected slowly using the micromanipulator. During the slow islet administration, delivery fluid was allowed to leak out from the hole where the micro-manipulator was inserted, leaving the islets in the anterior chamber of the eye. The pipette was removed and Refresh Tears supplemented with gentamicin (0.3% wt/vol) was applied to the eye to prevent desiccation and infection. Following surgery, the mice were returned to their cages on a warming pad (37$^{o}$C) (#E12107-919, Sunbeam) and monitored for recovery.

**Cytotoxicity Assays**

PBMCs from HLA-A2$^{+}$ and HLA-A2$^{-}$ donors were labelled with cell proliferation dye eFluor™ 450 (Thermo Fisher Scientific, #65-0842-85) and plated in 96 well U-bottom plates at 1 x $10^{6}$ cells per well. A2-CAR CD4$^{+}$ and CD8$^{+}$ T cells were added at the following A2-CAR T cells: PBMC ratios: 2:1, 1:2 and 1:10. Cultures were incubated for 24 hours, then stained with antibodies (**Supplementary Table S1**). To stain for caspase 3, cells were fixed using 2% formaldehyde and then permeabilized using Saponin.

**Flow Cytometry**

Analyses of cell phenotype and viability were performed after thawing of PBMCs, T cell enrichment and NGFR selection, as well as prior to A2-CAR T cell freezing. Antibodies used are listed in **Supplementary Table 1**. To quantify cell viability, apoptosis assays were conducted using the Apoptosis/Necrosis Assay kit (Abcam, #ab176749) in which cells are stained with 7-AAD, cytocalcein violet (CCV), and apopxin green[1]. Only cells which were >60% live, apoptosis-negative were used for in vivo injection.

To assess human cell engraftment in mice, blood was collected from the saphenous vein on a weekly basis. Red blood cells were lysed using RBC Lysis Buffer (Thermo Fisher Scientific, #00-4300-54), and cells were incubated with Rat Anti-Mouse CD16/CD32 (Mouse BD Fc Block™; BD, #553142). Cells were examined for viability and phenotyped using antibodies as listed in Table 1. 123count eBeads™ Counting Beads (Thermo Fisher Scientific, #01-1234-42) were added just prior to cytometer read out in order to quantify the number of cells in a given blood sample.

**Histology**

Kidneys with islet grafts were fixed in 4% PFA (Sigma Aldrich, #252449) and eyes with islet grafts were fixed in Davidson's fixative (two parts 37% formalin (Sigma Aldrich, #252549), three parts 100% ethanol (Commercial Alcohols, #P016EAAN), one part glacial acetic acid (Fisher, #A35-500) and three parts tap water) for 24-48 hours and stored in 70% ethanol (Commercial Alcohols, #P016EAAN) at

4°C until paraffin embedding. Paraffin-embedded blocks were cut into 5 µm-thick sections (Kidney capsule grafts: Wax-It Histology Services, Vancouver, Canada; ACE grafts: BCCHR Histology Core facility, Vancouver, Canada). Some slides were used for hematoxylin and eosin staining (Wax-it Histology Services, Vancouver, Canada; BCCHR Histology Core facility, Vancouver, Canada) and others were deparaffinized, rehydrated in graded ethanol, and immunostained as described[2]. Fluorescence images were captured with an ImageXpress® Micro XLS System (Molecular Devices Corporation; Sunnyvale, CA). For paraffin-embedded eye sections, islet images were captured using an Olympus BX61 Fluorescence and Bright Field Automated Upright Microscope with QImaging Retiga Exi camera and Olympus DP71 color camera. Primary and secondary antibodies are listed in Supplementary Table S1.

**Statistical Analyses**

Statistical analyses were performed in R version 4.1.2[3], using the tidyverse package[4]. Missing values for human C-peptide (below the limit of detection) were multiply imputed using Amelia[5] with an m parameter of the highest percentage of missing data for that dataset (10-20). Survival analyses were performed using packages survival and survminer[6,7] with pairwise comparisons using the Log-Rank test and corrections for multiple comparisons performed using the Benjamini-Hochberg false discovery rate method. For Figure 1F-I, multivariate regressions were performed using the lm() function from base R, with corrections for multiple comparisons performed using TukeyHSD(); an adjusted p-value of 0.05 was considered to be statistically significant and a null hypothesis of no difference. For **Figure 1H-I**, after visual inspection using qq plots for normal distribution, one-sample t tests were performed, comparing to the pre-injection level of 50%. Corrections for multiple comparisons were performed using the Benjamini-Hochberg method. For **Figure 3G-J**, a Bayesian approach with multilevel regression modelling was used due to the high number of correlated experiments with many repeated measurements[8], as well as non-normal distribution of the residuals (distributions used were skew normal, lognormal, or zero-inflated poisson). For each test, a mixed effects model was fitted with weeks post-A2-CAR T cell injection treated as categorical. If no result was reported, the posterior probability did not exceed 95% or the absolute difference was less than 10% (low biological significance). All Bayesian models were created in Stan computational framework accessed with the brms package[9,10]. Models were assessed and parameters adjusted to improve fit and/or address issues such as divergence, including treating individual animals as random variables contributing to Weeks, and adjusting the response distribution and link function. Models were compared using the leave-one-out methodology. In **Supplementary Figure S6 and S7**, lines are the fitted values of the posterior distribution with shading indicating the 50, 80, and 95% credible intervals. Related figures were generated using ggplot2[4]. See https://github.com/caraee/A2_CAR for scripts for performing the analyses and generating related figures.

**Supplemental Table 1.** *Antibodies used in the study.*

| Purpose | Target | Clone | Fluorophore | Company | Catalogue # |
|---|---|---|---|---|---|
| **In vitro phenotyping of expanded cells** | CD3 | UCHT1 | BV786 | BD Horizon | 564491 |
| | CD4 | OKT4 | BV510 | BioLegend | 317444 |
| | CD8a | RPA-T8 | PE-Cy7 | Invitrogen | 25-0087-42 |
| | CD14 | M5E2 | BV421 | Biolegend | 301830 |
| | CD271 (NGFR) | C40-157 | PE | BD Pharmingen | 557196 |
| | Myc | 9E10 | AF647 | UBC Ablabs | |
| | HLA-A*02 tetramer | N/A | APC | NIH Tetramer Facility | |
| | Fixable Viability Dye | N/A | eFluor™ 780 | Thermo Fisher Scientific | #65-0865-18 |
| **Ex vivo Phenotyping from blood** | hCD45 (anti human) | HI30 | V500 | BD Horizon™ | 560777 |
| | CD4 | SK3 | BV786 | BD Horizon™ | 563877 |
| | CD8a | SK1 | eF450 | eBioscience™ | 48-0087-42 |
| | CD271 | C40-157 | PE | BD Pharmingen™ | 557196 |
| | Myc | 9E10 | AF647 | UBC Ablabs | |
| | CD45 (anti mouse) | 30-F11 | PE-Cy7 | eBioscience | 25-0451-82 |
| **Cytotoxicity Assay** | CD3 | UCHT1 | BV786 | BD Biosciences | 564491 |
| | Active Caspase-3 | FITC | C92-605 | BD Pharmingen™ | 559341 |
| **Apoptosis Assay** | Apopxin Green Indicator | N/A | Ex/Em = 490/525 nm | abcam | ab176749 |
| | 7-AAD | N/A | Ex/Em = 546/647 nm | abcam | ab176749 |
| | CytoCalcein Violet 450 | N/A | Ex/Em = 405/450 nm | abcam | ab176749 |
| **Immunostaining of paraffin embedded tissue** | Insulin | Polyclonal guinea pig /IgG | NA | Invitrogen | PA1-26938 |
| | hCD45 | Mouse IgG1 clone H130 | NA | eBioscience | 17-0459-42 |
| | Anti-mouse Ab | Goat anti-mouse IgG (H+L) | APC | Invitrogen | A865 |
| | Anti-guinea pig Ab | Goat anti guinea pig IgG (H+L) | Alexa Flour 488 | Invitrogen | A11073 |

**Supplemental Table 2.** *Human Islet Donor Characteristics.*

| Donor ID | Sex | Age | HbA1c | HLA-A | Relevant Figs |
|---|---|---|---|---|---|
| R310 | Male | 25 | 5.4 | 02, 31 | Figs 1&2 |
| R330 | Female | 39 | 5.1 | 01, 02 | Figs 3&4 |
| R427 | Male | 52 | 5.8 | 01, 23 | Fig 5 |
| R439 | Female | 57 | NA | 02, 26 | Fig 5 |

NA = not available

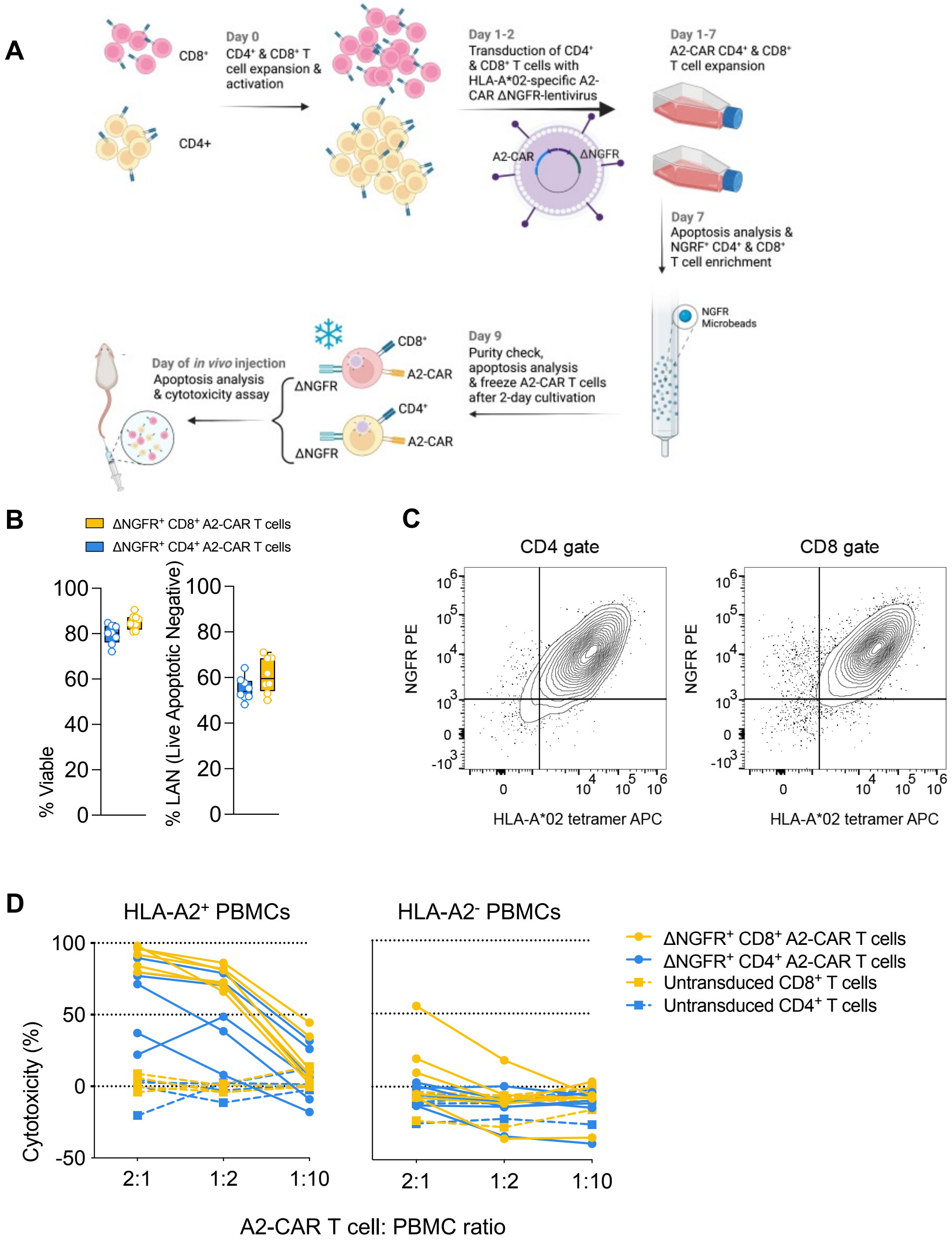


**Figure S1.** *In vitro* generation and characterization of human A2-CAR T-Cells prior to *in vivo* studies. **(A)** Overview of A2-CAR T cell generation process. **(B)** Viability of ΔNGFR-purified A2-CAR CD4$^+$ and CD8$^+$ T cells prior to mouse injection. Left: viability data reported by apoptosis assay. Right: % Live, apoptosis negative (LAN). Boxplot; Minima: minimum outliers; Maxima: maximum outliers; Centre: median; Bounds of box: first to third quartile; Whiskers: The whiskers extend from the smallest value up to the largest value. **(C)** Representative flow cytometry plots showing coordinate expression of DNGFR and the A2-CAR detected by staining with an HLA-A2 tetramer. **(D)** Cytoxicity capacity of untransduced and A2-CAR-transduced CD4$^+$ and CD8$^+$ T cells (n=6), towards human HLA-0*2 positive and negative PBMCs. Data from 4 independent experiments.

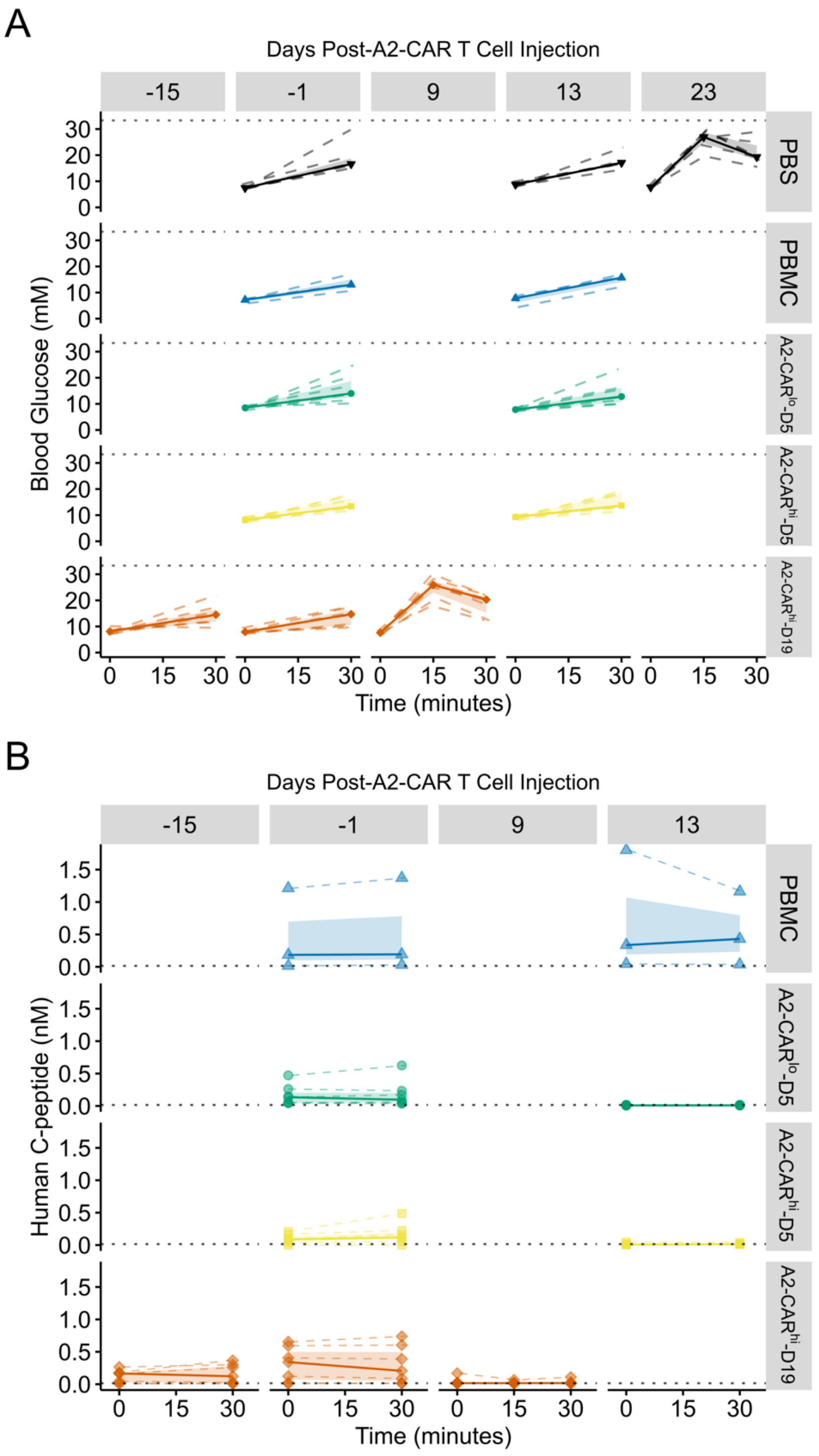


**Figure S2.** *Additional glucose stimulated insulin secretion data.* Related to Figure 1. Animals were given 2 g/kg of glucose intraperitoneally after a 6 hour fast, starting between 0730 and 0900. Blood was collected from the saphenous vein at and 30 minutes after glucose injection, except for the A2-CAR$^{hi}$-D19 group, from which blood was also collected at 15 minutes post-glucose injection. **(A)** Blood glucose and **(B)** plasma human C-peptide were measured. Solid lines are the medians, shading is the interquartile range, and dashed lines are individual mice. Dotted horizontal lines indicated the limit of detection of blood glucose (33.3 mM) **(A)** or human C-peptide (7.4 pM) **(B)**.

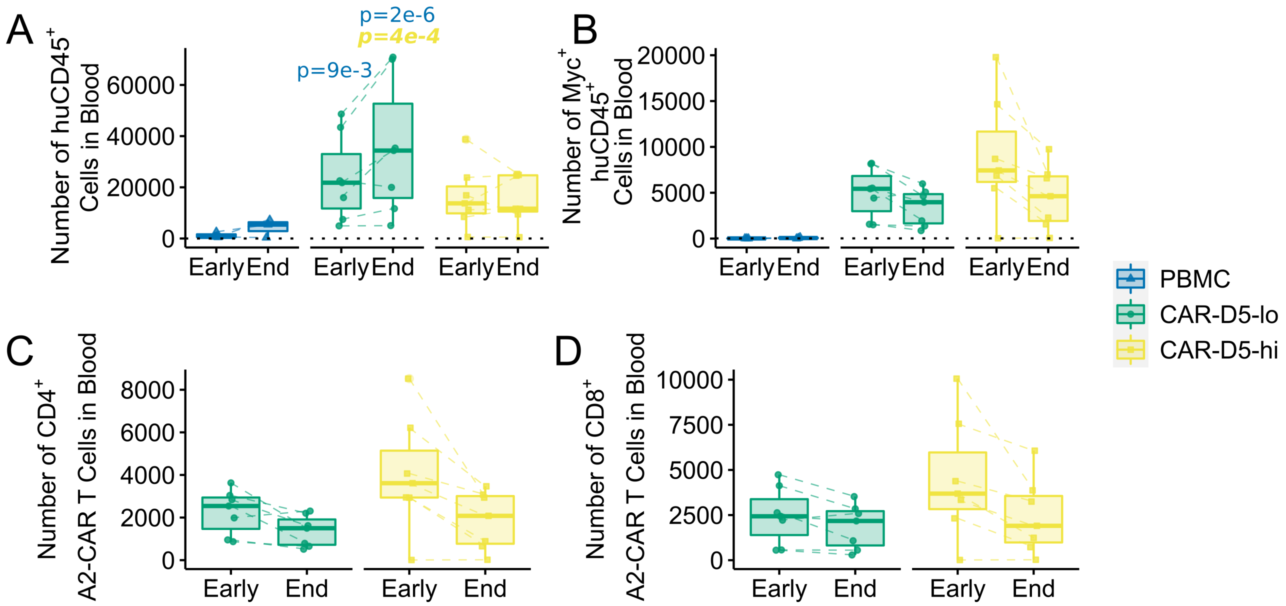


**Figure S3.** *Immune cell absolute counts.* Related to Figure 1. The absolute number of the indicated T cells was measured in blood on day 9 or 13 (Early) or 16 (End). **(A)** human CD45$^+$ and **(B)** A2-CAR (Myc$^+$) T cells. Within the A2-CAR$^+$ population, the numbers of **(C)** CD4$^+$ and **(D)** CD8$^+$ T cells was measured. **(A-B)** Dotted horizontal lines indicates the limit of detection of human CD45+ cells. Colour and style of the p-values indicates direction of comparison: regular indicates a comparison to PBMC, and ***bold italics*** indicates a comparison to A2-CAR$^{hi}$-D5.

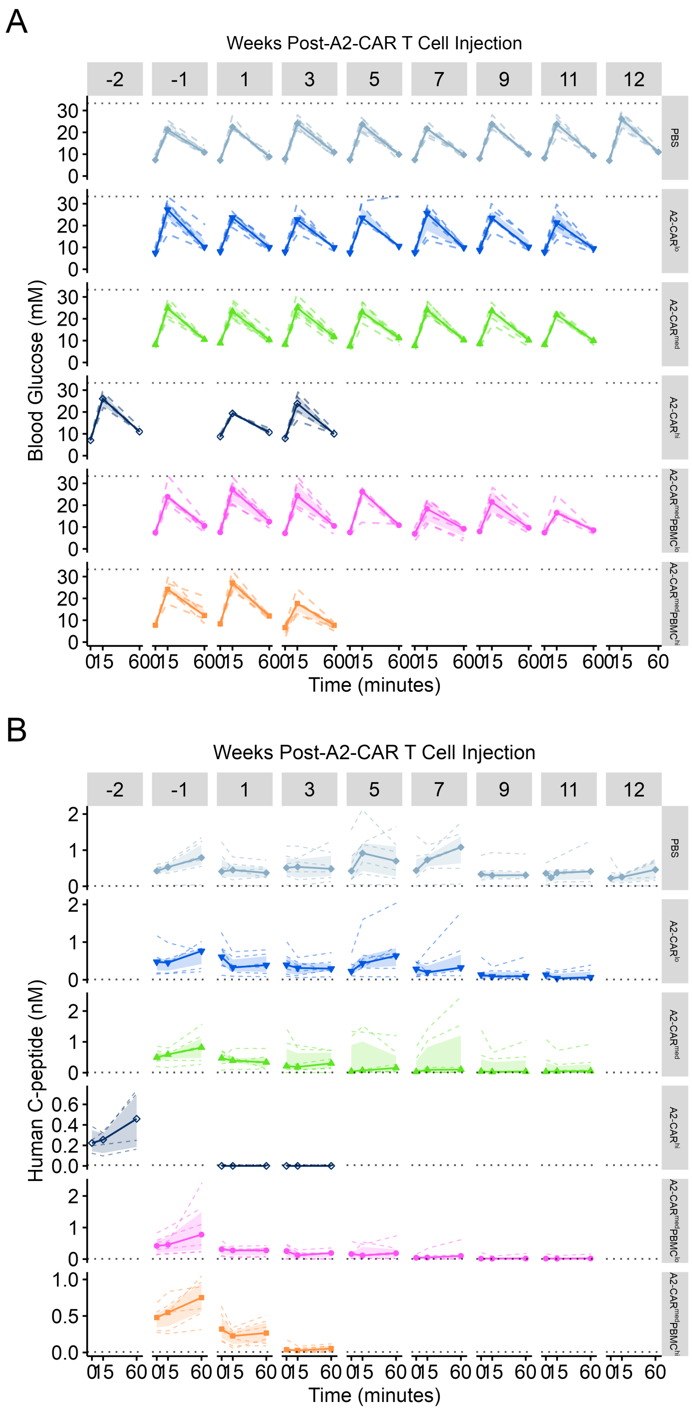


**Figure S4.** *Additional glucose stimulated insulin secretion data.* Related to Figure 3. Animals were given 2 g/kg of glucose intraperitoneally after a 6 hour fast, starting between 0730 and 0900. Blood was collected from the saphenous vein before and 15 and 60 minutes after glucose injection. **(A)** Blood glucose and **(B)** plasma human C-peptide were measured. Solid lines are the medians with points indicating the times of measurements, shading is the interquartile range, and dashed lines are individual mice. Dotted horizontal lines indicated the limit of detection of blood glucose (33.3 mM) **(A)** or human C-peptide (14.9 pM) **(B)**. The week 12 data for the PBS group are the same data as for week -2 in the A2-CAR$^{hi}$ group as a subset of the PBS-treated mice were injected with 3 million A2-CAR T cells at 18 weeks post-islet transplantation.

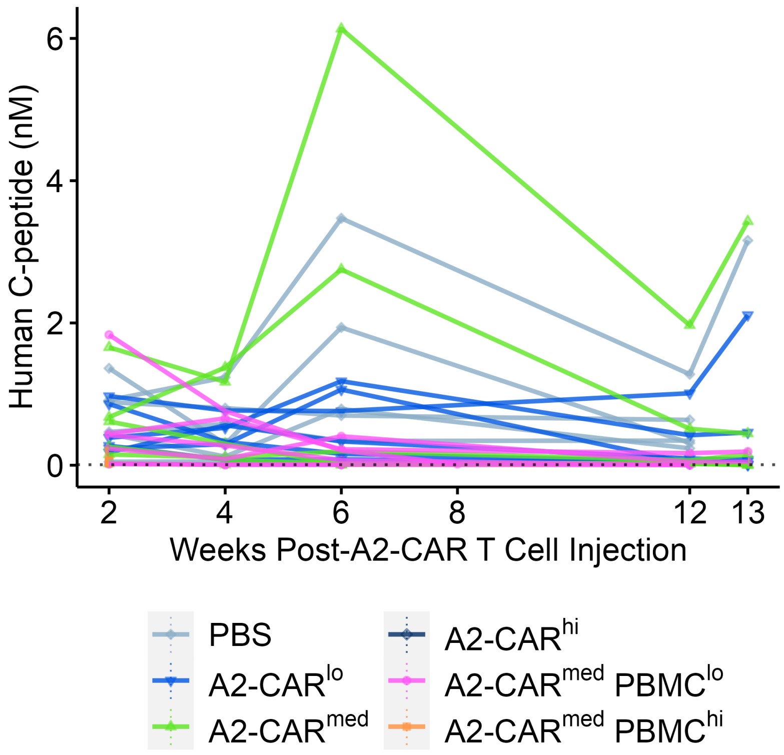


**Figure S5.** *Random fed human C-peptide measurements.* Related to Figure 3. Blood was collected from the saphenous biweekly in the morning (without fasting) and assayed for plasma human C-peptide. Points and solid are individual mice. Dotted horizontal lines indicated the limit of detection of human C-peptide (14.9 pM).

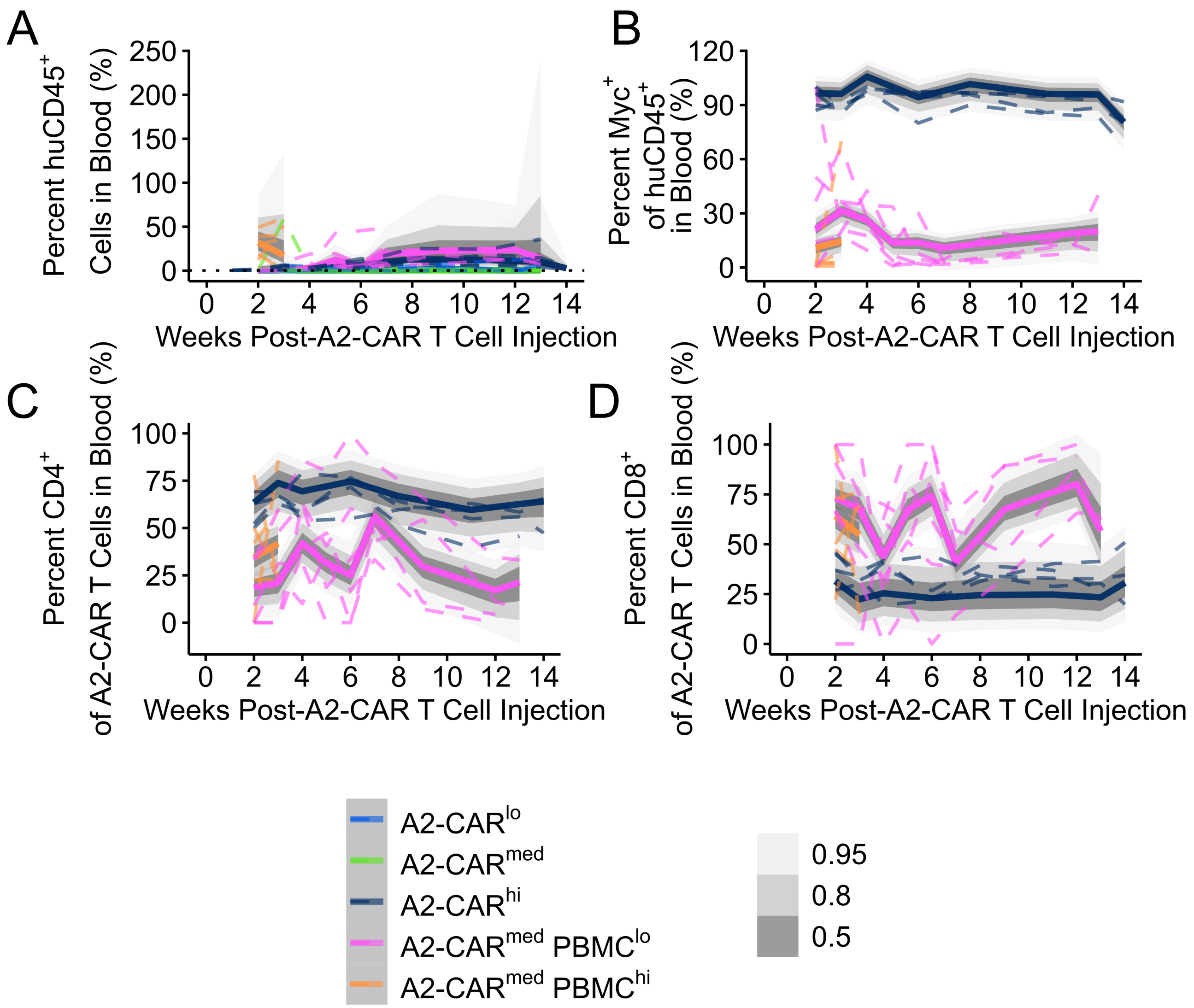


**Figure S6**. *Bayesian mixed effects models for human immune cell engraftment by percentage.* Related to Figure 3. Blood was collected from the saphenous weekly post-A2-CAR-T cell injection and flow cytometry was used to determine the percentages of **(A)** human CD45$^{+}$ and **(B)** Myc$^{+}$ cells. Within the Myc+ gate, the proportions of **(C)** CD4$^{+}$ and **(D)** CD8$^{+}$ cells were measured. Bayesian models were used to fit to the data for statistical analyses. In all panels dashed lines are individual animals, solid lines are the mean of 1000 draws from each of the four chains of the posterior predictive distribution for each model, and shading indicates the predictive bands at the 50%, 80%, and 95% levels. Evidence ratios are shown in **Figure 3**, but where the bands do not overlap the evidence ratio is at least 20.

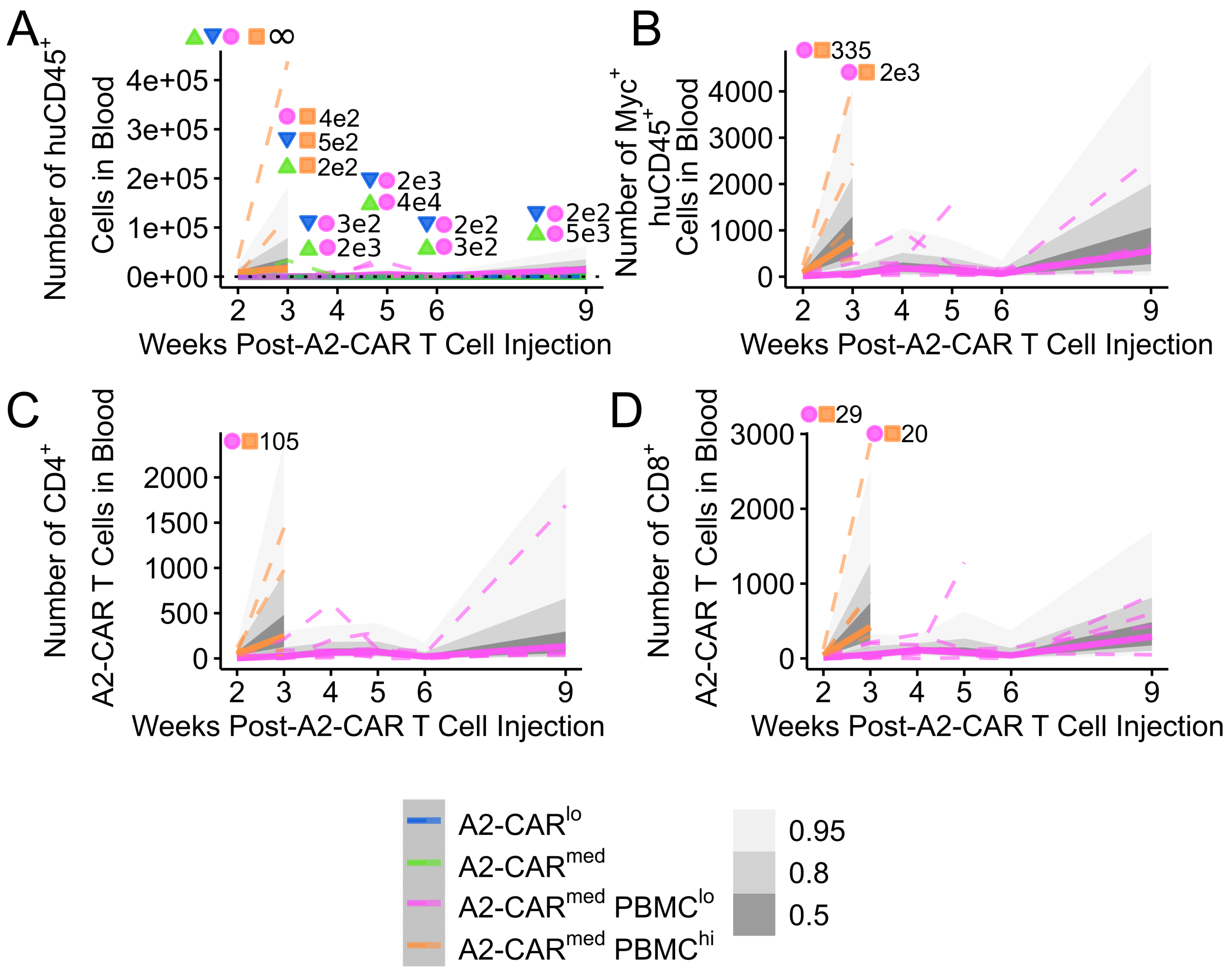


**Figure S7**. *Bayesian mixed effects models for human immune cell engraftment by cell count.* Related to Figure 3. Blood was collected from the saphenous weekly post-A2-CAR-T cell injection, count beads were added, and the number of **(A)** human CD45$^{+}$ and **(B)** Myc$^{+}$ cells was measured. Within the Myc+ gate, the proportions of **(C)** CD4$^{+}$ and **(D)** CD8$^{+}$ cells were measured Bayesian models were fit to the data for statistical analyses. In all panels, dashed lines are individual animals, solid lines are the mean of 1000 draws from each of the four chains of the posterior predictive distribution for each model, and shading indicates the predictive bands at the 50%, 80%, and 95% levels. Values are Bayes Factors and symbols show comparisons.

### References

1. Fung VCW, Rosado-Sanchez I, Levings MK. Transduction of Human T Cell Subsets with Lentivirus. *Methods Mol Biol.* 2021;2285: 227-254.

2. Asadi A, Bruin JE, Kieffer TJ. Characterization of Antibodies to Products of Proinsulin Processing Using Immunofluorescence Staining of Pancreas in Multiple Species. *J Histochem Cytochem.* 2015;63(8): 646-662.

3. *R: A language and environment for statistical computing* [computer program]. Vienna2022.

4. Wickham H. *ggplot2: Elegant Graphics for Data Analysis.* New York: Springer-Verlag; 2016.

5. Honaker J, King G, Blackwell M. Amelia II: A Program for Missing Data. *Journal of Statistical Software.* 2011;45: 7.

6. *survminer: Drawing Survival Curves using 'ggplot2'_. R package version 0.4.9* [computer program]. 2021.

7. *A Package for Survival Analysis in R_. R package version 3.3-1* [computer program]. 2022.

8. Gelman A, Hill J, Yajima M. Why We (Usually) Don't Have to Worry About Multiple Comparisons. *Journal of Research on Educational Effectiveness.* 2012;5(2): 189-211.

9. Bürkner P-C. brms: An R Package for Bayesian Multilevel Models Using Stan. *Journal of Statistical Software.* 2017;80(1): 1-28.

10. Bürkner P-C. Advanced Bayesian Multilevel Modeling with the R Package brms. *The R Journal.* 2018;10(1): 395-411.
